# Publishing of COVID-19 Preprints in Peer-reviewed Journals, Preprinting Trends, Public Discussion and Quality Issues

**DOI:** 10.1101/2020.11.23.394577

**Authors:** Ivan Kodvanj, Jan Homolak, Davor Virag, Vladimir Trkulja

**Affiliations:** Department of Pharmacology, University of Zagreb School of Medicine, Zagreb, Croatia

**Keywords:** preprint, COVID19, peer-review, publishing

## Abstract

**Introduction:** COVID-19-related (vs. non-related) articles appear to be more expeditiously processed and published in peer-reviewed journals. We aimed to evaluate: (i) whether COVID-19-related preprints were favored for publication, (ii) preprinting trends and public discussion of the preprints, and (iii) the relationship between the publication topic (COVID-19-related or not) and quality issues.

**Methods:** Manuscripts deposited at bioRxiv and medRxiv between January 1 and September 27 were assessed for the probability of publishing in peer-reviewed journals, and those published were evaluated for submission-to-acceptance time. The extent of public discussion was assessed based on Altmetric and Disqus data. The Retraction Watch Database and PubMed were used to explore the retraction of COVID-19 and non-COVID-19 articles and preprints.

**Results:** With adjustment for the preprinting server and number of deposited versions, COVID-19-related preprints were more likely to be published within 120 days since the deposition of the first version (OR=2.04, 95%CI 1.87-2.23) as well as over the entire observed period (OR=1.42, 95%CI 1.33-1.52). Submission-to-acceptance was by 38.67 days (95%CI 34.96-42.39) shorter for COVID-19 articles. Public discussion of preprints was modest and COVID-19 articles were overrepresented in the pool of retracted articles in 2020.

**Conclusion:** Current data suggest a preference for publication of COVID-19-related preprints over the observed period.

## 1. INTRODUCTION

The concepts of preprinting, a practice of presenting scientific/scholarly papers to the scientific community before they are published in peer-reviewed journals, date back to the 17^th^ century (Moore, 1965). In the modern era, one of the first large-scale attempts to share unpublished works was the Information Exchange Group (IEG) experiment. It was carried out by the National Institute of Health (NIH) in the 1960s (Cobb, 2017), with the goal of speeding up information dissemination and promoting discussion (Green, 1964). Although the project had numerous members, it was brought to an early and infamous end after several discouraging editorials had been published (Abelson, 1966; “Four years of information exchange,” 1966). The proposal for the Physics Information Exchange (PIE) (Cobb, 2017) elicited similar unfavorable reactions. It was called offensive (“Unpublished Literature,” 1966), and declared “a serious threat to physics communication and to the physics research community” by the editor of the Physical Review (Pasternack, 1966). After this setback, it took years for PIE to start, but finally, it evolved into the first major preprint server (Till, 2001) – arXiv - currently distributing articles in the fields of physics, mathematics, computer science, quantitative biology, quantitative finance, statistics, electrical engineering and systems science, and economics (“arXiv,” n.d.).

While preprinting is a standard practice in some fields (Berg et al., 2016), it is still uncommon in biomedicine (Fu and Hughey, 2019). Nowadays this seems like a paradox since the IEG, one of the first attempts to establish a platform for sharing preprints, was focused on the biomedical field. It took years for biomedical preprints servers to be established, even though the initiative was present but often disregarded. For instance, Harold Varmus, Nobel prize winner and director of NIH, proposed the development of a preprint repository to “accelerate much-needed public discussion of electronic publication” (Varmus, 1999) in 1999. After several failed attempts to establish preprints servers (Nature Precedings and ClinMed Netprints) (“About Precedings,” n.d.; Cobb, 2017), bioRxiv was finally started (Callaway and Powell, 2016) - currently one of the largest preprint servers in the field.

Through preprints, authors can rapidly share their findings and receive feedback from other scientists. Both sides can profit from this - authors by improving their manuscripts before submitting them to journals, based on the received feedback; whilst others could steer their research in accordance with the newest findings that would otherwise be unknown for months. Furthermore, some authors even believe that the lack of peer-review is beneficial since it does not restrain the sharing of worthwhile ideas that might be falsely disregarded and overlooked in the editorial process (Green, 1964). On the other hand, there is a concern that without peer-review, preprints might be of lower quality (Nabavi Nouri et al., 2020). Both bioRxiv and medRxiv explicitly declare that preprints “should not be regarded as conclusive, guide clinical practice/health-related behavior, or be reported in news media as established information”. However, peer-review is also prone to bias and the evidence of its effectiveness is scarce (Carneiro et al., 2020; Jefferson et al., 2002; Smith, 2006; Vercellini et al., 2016).

The Coronavirus disease 2019 (COVID-19) pandemic prompted a need for rapid communication and discussion of scientific information and resulted in an unprecedented surge in the number of publications on various aspects of the disease. Publishers greatly shortened the peer-review process for COVID-19-related manuscripts (Homolak et al., 2020; Horbach, 2020; Kun, 2020). Be it due to elementary greed or to a praiseworthy desire to disseminate information, such a practice is not necessarily beneficial – concerns about the quality of articles published under these circumstances have been raised (Homolak et al., 2020). Preprinting apparently became a necessity – not only by enabling rapid dissemination of information but also by encouraging public discussion and providing an opportunity for peers to intervene early if methodological fallacies are identified and limit the spread of misinformation to improve the overall quality of the manuscripts. We aimed: a) to evaluate preprinting trends for COVID-19-related and non-related manuscripts on two key biomedical preprint platforms *bioRxiv* and *medRxiv* during the COVID-19 pandemic; b) to estimate the extent to which the preprints were commented on (as an indicator of the extent of “pre-submission peer-review”); c) to estimate whether COVID-19 manuscripts are favored over non-COVID-19 manuscripts for journal publishing; d) to probe the feasibility of the assumption that published COVID-19 papers are (under the circumstances of shortened peer-review process) more commonly flawed than the published non-COVID-19 papers.

## 2. MATERIALS AND METHODS

### 2.1. Study outline and study outcomes

Types of datasets used and their purpose in the present study are outlined in Figure 1. We defined three objectives based on their relevance (objective 1 – most relevant, objective 3 – least relevant) and the anticipated level of susceptibility to bias/confounding (objective 1 – least susceptible, objective 3 – most susceptible), given the nature of the study and the type of data. The 1^st^ objective was to investigate whether COVID-19-related preprints were favored (over non-related) for publication in peer-reviewed journals. We considered that the probability of publishing was the most informative outcome in this respect and that further insight would be obtained from submission-to-acceptance data. Therefore, we defined three outcomes for the purpose: (i) *primary outcome* was the probability of publishing within 120 days since the deposition of the first preprint version and was assessed in a subset of preprints deposited till June 29, 2020. Since the time-period during which manuscripts could have been preprinted and published was bounded (January 1 – November 01, 2020), to reduce the risk of bias arising from unequal “time-at-risk” (for publishing), the analysis was restricted to preprints deposited till June 29 and a time-window for the publishing of 120 days. We considered it to be a reasonable period of time for the submission and review process to take place, and it was “available” to all preprints in this subset; (ii) *secondary outcome* was the probability of publishing over the entire observed period (i.e., before November 01, 2020) and it was assessed in a subset of preprints deposited before September 27, 2020. All articles published after November 01, 2020, were considered unpublished. We considered this outcome to be complementary to the primary outcome, which was susceptible to bias arising from a possibility that not all preprinted manuscripts were actually submitted to journals at the same time, i.e., that they might have been purposely left in a preprint form over a longer period of time. To further reduce bias/confounding arising from unequal “time-at-risk”, preprints in both datasets were stratified into 15-day strata regarding the date of the first preprinted version. Due to the limited number of COVID-19 related preprints, the first two strata (Jan. 01 - 15 and Jan. 16 - 30) were merged into one stratum; (iii) *tertiary outcome* was submission-to-acceptance time, considered a proxy of the length of the peer-review process, and was assessed in a subset of preprints that were published in peer-reviewed journals during the observed period and submitted to journals after January 01, 2020. Here, we anticipated potential confounding arising from varying interest in COVID-19-related and non-related topics over time, so manuscripts were stratified into 15-day strata regarding the journal submission dates. The 2^nd^ objective was to illustrate preprinting trends of COVID-19-related and non-related manuscripts on bioRxiv and medRxiv, their usage statistics and to estimate the extent of the public peer-review (i.e., pre-submission peer-review) using the number of posted comments and Altmetric data as proxies. This was assessed using all preprints deposited at the two platforms between January 1 and December 05, 2020. The 3^rd^ objective was to evaluate a possible association between the publication topic (COVID-19-related or non-related) and quality issues related to the published papers using notifications on retraction or issuance of concerns or corrections as proxies. This was assessed using data on all published papers indexed in PubMed and all notifications issued in the Retraction Watch Database (“Retraction Watch Database,” n.d.) between January 1 and December 05, 2020.

**Figure 1.**
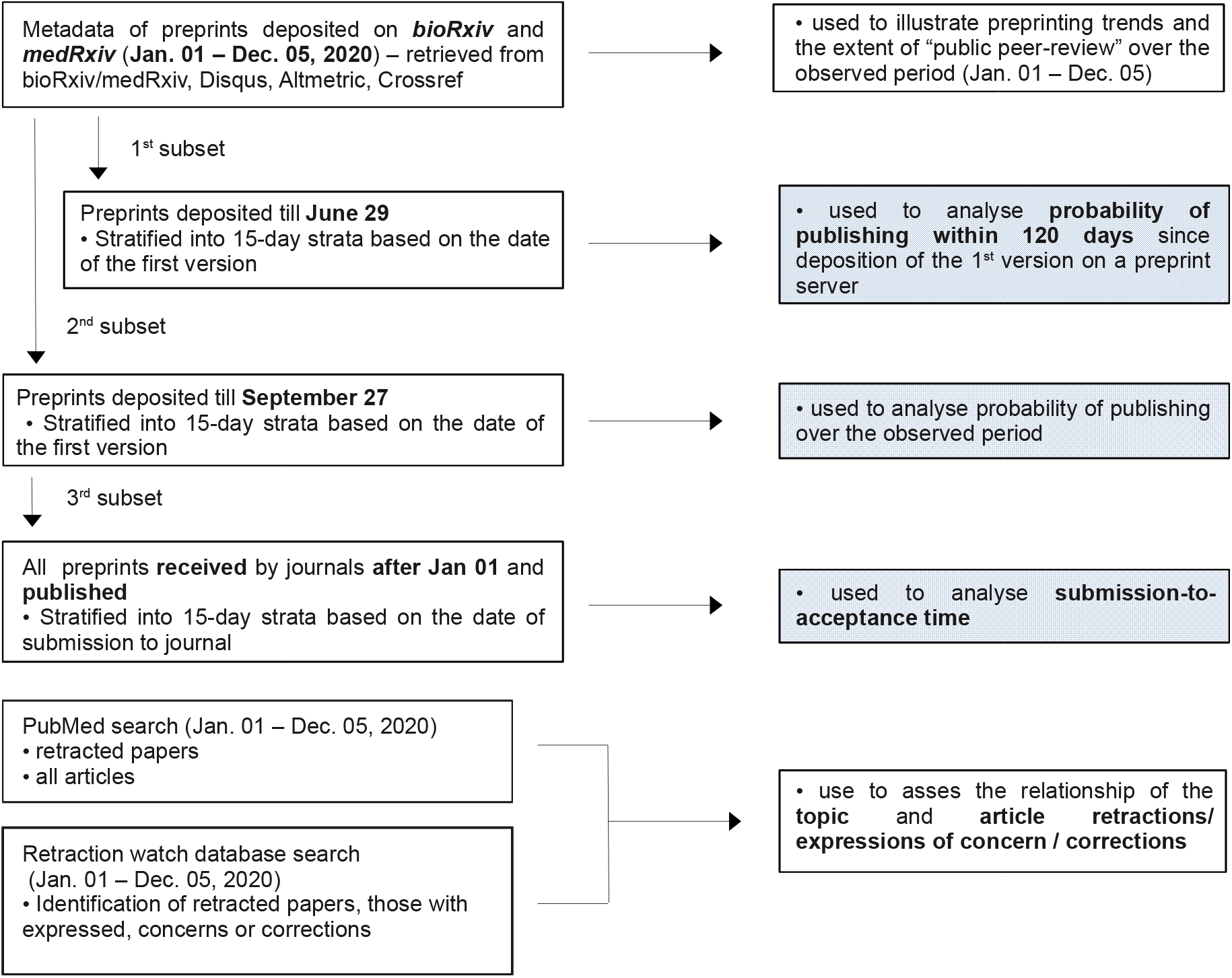
Study outline. Measure outcomes of the primary objective are marked in blue.

### 2.2. Data collection

All data retrieval and management were done in *R* (version 4.0.2) (“The R Project for Statistical Computing,” n.d.). *Rbiorxiv* and *medrxivr* packages were used to access bioRxiv and medRxiv application programming interfaces (APIs) and to collect metadata and usage statistics on the preprinted manuscripts (January 1 – December 05, 2020). COVID-19 preprints were identified using the search terms COVID-19 OR SARS-CoV-2 OR Coronavirus disease 19 OR 2019-nCoV. All other preprints were classified as non-COVID-19 preprints. Publication status was retrieved from bioRxiv and medRxiv services. Publication dates of articles with bioRxiv preprints were provided by the bioRxiv API. *Rcrossref* was used to gather publication dates for journal-published preprints, initially deposited on the *medRxiv*. The *DisqusR* and *rAltmetric* packages were used to access the Disqus and Altmetric API and identify the number of comments and to retrieve Altmetric data for each preprint deposited on bioRxiv and medRxiv servers. Submission and acceptance dates for published preprints were fetched from PubMed with the *RISmed* package. The Retraction Watch Database was used to retrieve the number of COVID-19-related articles with issued retraction notice, expression of concern, or correction during 2020 (till December 05). To retrieve “control groups”, 7 search phrases were constructed: 4 were used to retrieve the number of retraction notice, expression of concern, or correction pertaining to chosen viruses and their associated diseases (HIV, Hepatitis virus, Herpes virus and Influenza); 2 were used to retrieve the numbers pertaining to 2 topics (immunology and epidemiology of viral infectious diseases); 1 was used to retrieve the number of retraction notice, expression of concern or corrections issued for all COVID-19-unrelated articles (total number - COVID-19 number). The exact search methodology is depicted under Table 3. All numbers are expressed as a proportion of the total number of articles indexed by PubMed. To minimize the time-at-risk risk bias, only a comparison of articles published in 2020 was made. Albeit, the risk of bias still exists due to the possibility of different publishing rates of articles during 2020. To avoid bias due to possible over- or underrepresentation of the COVID-19 articles in PubMed, in comparison with the Retraction Watch database, we repeated the analysis using only data provided by the PubMed database. The search was conducted using the search term (Retracted Publication[PT] OR “Retraction:”[title]) AND (“2020/01/01”[Date - Publication] : “2020/12/05”[Date - Publication]), with and without “AND (COVID-19 OR SARS-CoV-2 OR “Coronavirus disease 19” OR 2019-nCoV)”, to identify the number of retractions of COVID-19 and non-COVID-19 articles during 2020.

**Table 3.** The number of retractions/expressions of concern/corrections identified at Retraction Watch Database (RWD), the total number of PubMed articles, and retraction rates (notices/total number of articles) for COVID-19-related articles and articles related to four other viruses/associated diseases, two research fields (epidemiology, immunology).

| *Topic* | *RWD count<sup>1</sup>* | *Pubmed count<sup>2</sup>* | *Retraction rate* |
| --- | --- | --- | --- |
| COVID-19 | 36 | 64915 | 0.55‰ |
| Not COVID-19 | 306 | 1428252 | 0.21‰ |
| HIV | 1 | 10453 | 0.10‰ |
| Herpes virus | 3 | 2663 | 1.13‰ |
| Influenza | 0 | 3411 | - |
| Hepatitis virus | 0 | 1321 | - |
| Epidemiology | 6 | 40279 | 0.15‰ |
| Immunology | 8 | 63577 | 0.13‰ |

<sup>1</sup> Search phrases used for searching Retraction Watch Database: COVID-19 OR SARS-CoV-2 OR Coronavirus disease 19 OR 2019-nCoV; HIV OR AIDS OR human immunodeficiency virus; Herpes OR Herpes simplex OR HSV1 OR HSV2 OR varicella-zoster OR HZV OR Cytomegalovirus OR CMV OR Epstein-Barr OR EBV; Influenza OR Influenzavirus OR flu OR \*influenza\* OR H1N1 OR H5N1; Hepatitis virus OR HepA OR HepB OR HepC OR HepD OR HepE OR HAV OR HBV OR HCV OR HDV OR HEV OR Viral hepatitis; epidemiol\* OR infect\* OR virology OR virus OR viral; immun\* OR inflamma\* OR cytokine\* OR leukocy\* OR neutroph\* OR lymphoc\* OR antigen\* OR antibod\*. Search was limited to articles published from Jan. 01, 2020 till Dec. 05, 2020, with notice published during the same period. All preprints were omitted and all COVID-19 articles were excluded where the topic was not related to COVID-19.
<sup>2</sup> Same search phrases were used in conjunction with "AND ("2020/01/01"[Date - Publication] : "2020/12/05"[Date - Publication]) NOT (COVID-19 OR SARS-CoV-2 OR "Coronavirus disease 19" OR nCoV-19) NOT Preprint[Publication Type]", and with "[title]" added next to each term.

### 2.3. Data analysis

All data visualization and analysis were performed in *R* (version 4.0.2) (“The R Project for Statistical Computing,” n.d.). Data on preprinting trends over time, preprint usage statistics, Altmetric data and Disqus comments on the manuscripts preprinted during the observed period, and Retraction Watch Database data on the published papers were summarized by the preprinting platform and topic (COVID-19-related or not).

Probability of publishing within 120 days since the 1^st^ preprint version and the probability of publishing over the entire observed period were analysed by fitting stratified (in respect to preprint date) logistic regression (package *survival*, function clogit). Submission-to-acceptance time was analyzed by fitting a hierarchical (mixed) model with the submission date stratum as a random effect. Articles with submission dates before 2020 were excluded from the dataset that was used for analyzing submission-to-acceptance time. Fixed effects in all analyses were topic (COVID-19-related or non-related), preprinting platform (bioRxiv or medRxiv), and the number of preprinted versions (dichotomized as one or ≥2). The latter adjustment was introduced to account for potential bias arising from different intentions of the preprinting authors. For example, a preprint might have been submitted to a journal at the time of preprinting, or it might have been a work in progress with several versions and purposely (only) kept as a preprint over a longer period of time; more preprinted versions might have improved the quality of the final submitted version, hence peer-review process might have been shorter.

We conducted a supplemental analysis of preprinting/publishing trends and other aspects of bioRxiv and medRxiv preprints over a longer period of time that we found informative for discussion of the results of the main study (see Supplemental Material, Supplemental methods).

### 2.4. Submission-to-acceptance dataset validation

After all articles with submission dates before January 1, 2020, were removed, we identified and excluded one article with the submission-to-acceptance time of −1 days (an obvious mistake). To validate the dataset, submission and acceptance dates were checked for 100 randomly selected articles. For 1 article we could not find the submission and acceptance dates on the journal website, nor published pdf of the article. One article with erroneous values was identified (the acceptance date was off by 1 day). We considered that the number and size of errors were acceptable and not likely to affect the conclusions of the study.

## 3. RESULTS

### 3.1. Are COVID-19-related preprints favored for publishing in peer-reviewed journals?

The subset of preprints deposited till June 29, 2020, and used to evaluate the probability of publishing within 120 days since the 1^st^ preprint version comprised a total of 18810 preprints on bioRxiv (6.34% COVID-19-related and 93.66% non-related) and a total of 6576 preprints on medRxiv (64.92% COVID-19-related and 35.08% non-related) (Fig. 2). The subset of preprints deposited till September 27, 2020, and used to evaluate the probability of publishing over the entire observed period (before November 01, 2020) comprised 28481 preprints on bioRxiv (6.86% COVID-19-related and 93.14% non-related) and 10320 preprints on medRxiv (63.67% COVID-19-related and 36.33% non-related) (Fig. 2). Raw proportions of published papers by the topic and preprinting platform are depicted in Figure 2. In multivariate analyses (Table 1), COVID-19-related preprints were associated with higher odds of publishing within 120 days than non-COVID-19 preprints (OR=2.04, 95%CI 1.87-2.23), and with a higher probability of publishing over the observed period than non-COVID-19 preprints (OR=1.42, 95%CI 1.33-1.52). The probability for both outcomes was higher for preprints deposited on bioRxiv and lower for preprints with ≥2 versions than for those with only one version (Table 1). Journal submission-to-acceptance time was identified for preprints published before November 01, 2020. In multivariate analysis, the COVID-19-related topic was associated with almost 40 days shorter submission-to-acceptance time (mean difference −38.67, 95%CI −42.39 to −34.96) (Table 2). The preprinting platform and number of preprint versions did not appear associated with the outcome (Table 2).

**Table 1.** Summary of the analysis of the probability of publishing within 120 days since the first preprint version (a subset of preprints deposited till June 29) and time-to-publishing considering the entire observed period (a subset of preprints deposited till 27 September)^1^. Effects are expressed as odds ratios (OR).

| Predictors | Published within 120 days |  | Published before Nov. 01 |  |
| --- | --- | --- | --- | --- |
|  | OR (95% CI) | p | OR (95% CI) | p |
| COVID-19 vs Not COVID-19 | 2.04 (1.87 to 2.23) | **<0.001** | 1.42 (1.33 to 1.52) | **<0.001** |
| bioRxiv vs medRxiv | 1.33 (1.21 to 1.45) | **<0.001** | 1.24 (1.17 to 1.32) | **<0.001** |
| ≥2 vs 1 version | 0.72 (0.66 to 0.78) | **<0.001** | 0.90 (0.85 to 0.94) | **<0.001** |

<sup>1</sup> Stratified (by preprinting date of the first version) logistic regression was fitted to the probability of being published within 120 days and over the entire observed period.

**Table 2.** Analysis of *submission-to-acceptance* time for published preprints (in days)^1^.

| Predictors | Estimates (95% CI) | p |
| --- | --- | --- |
| COVID-19 vs Not COVID-19 | -38.67 (-42.39 to -34.96) | **<0.001** |
| bioRxiv vs medRxiv | 0.25 (-3.36 to 3.86) | 0.891 |
| ≥2 vs 1 preprint version | 0.96 (-2.00 to 3.91) | 0.526 |

<sup>1</sup> Linear mixed-effects model (submission date specified as a random effect) was fitted to the number of days between the date of receipt by the journal and date of acceptance. Submission to publication time was identified for 421 COVID-19/bioRxiv (median=49 days), 4191 not-COVID-19/bioRxiv (median=95 days), 1032 COVID-19/medRxiv (median=48 days), 423 not-COVID-19/medRxiv preprints (median=97 days).

**Figure 2.**
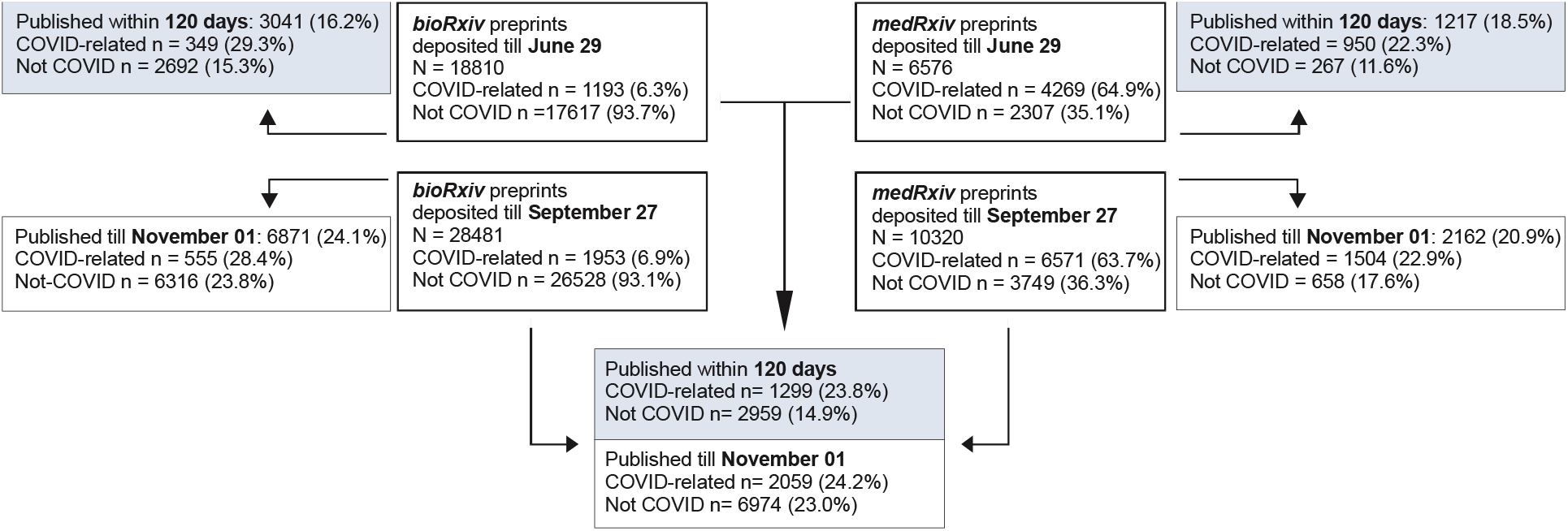
Structure of preprints deposited by June 29 and September 27, used to evaluate the probability of publishing by platform-by topic-by publishing outcomes. Blue color marks the primary measure outcome of the primary objective.

### 3.2. Preprinting trends, usage statistics, and indicators of public pre-submission peer-review

For the observed period (Jan. 1 – Dec. 05, 2020) we identified a total of 13257 preprints newly deposited at medRxiv [8298 (62.6%) COVID-19-related)], and 36267 preprints deposited at bioRxiv [2493 (6.87%) COVID-19-related]. There was a clear increasing trend in the number of newly deposited preprints (Fig 3A), but on medRxiv, the increase was largely due to the increasing number of COVID-19-related preprints, while the number of newly deposited preprints on bioRxiv appeared comparable for COVID-19-related and not related topics (Fig. 3A). Usage statistics of bioRxiv preprint server indicated an increase in the number of abstract views, full-text views, and PDF downloads (Fig. 3B).

**Figure 3.**
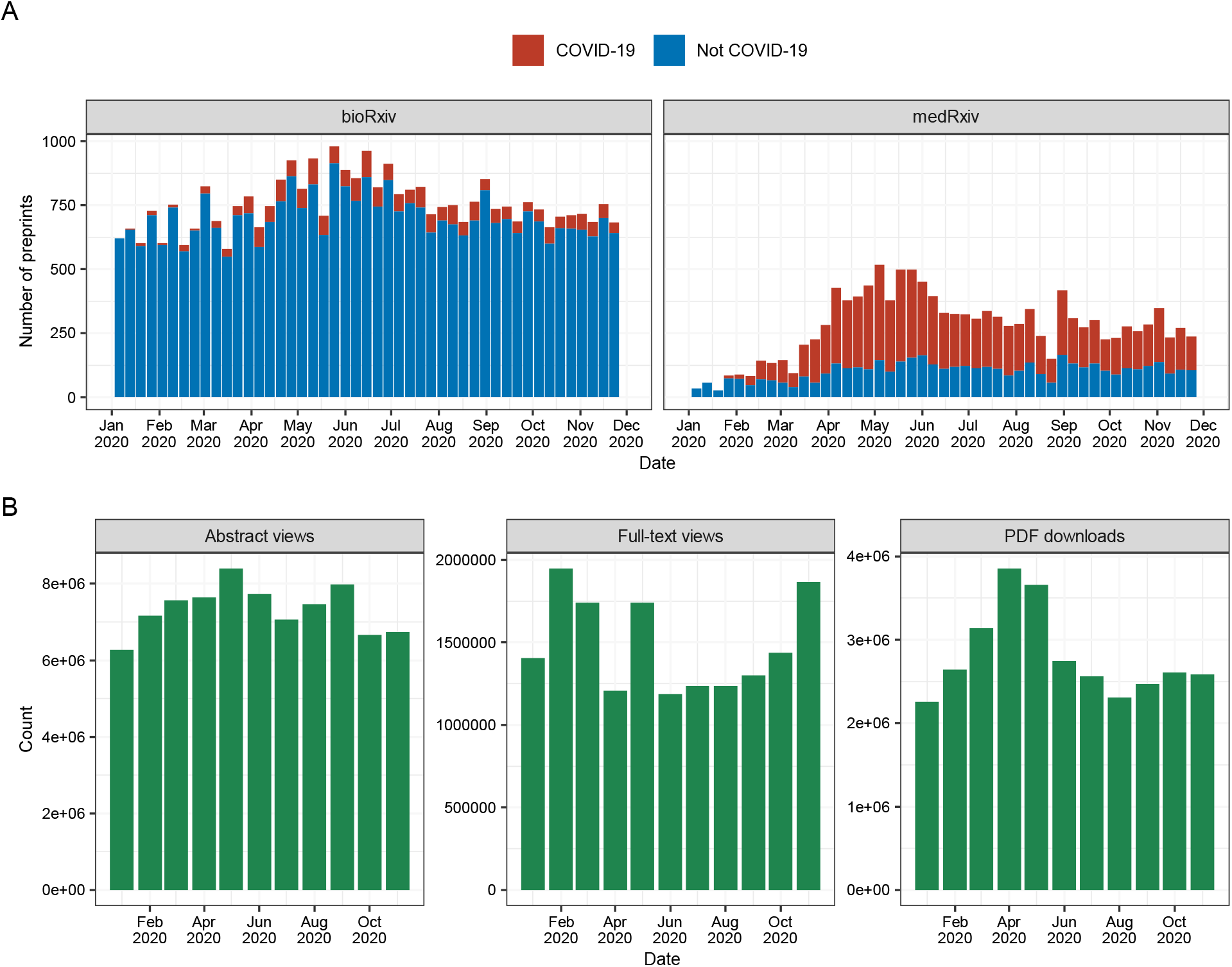
Posting of preprints and bioRxiv server usage statistics. **(A)** The number of new preprints posted on bioRxiv and medRxiv over the observed period. **(B)** Monthly abstract views, full-text views, and PDF downloads on bioRxiv server (not available for medRxiv).

The overall proportion of preprints that have been commented on is rather low (5.7%), but somewhat higher for COVID-19-related preprints (Fig. 4A): 21.6% and 3.2% of the COVID-19-related and not related preprints respectively, on bioRxiv; and 12.9% and 2.4% respectively, on medRxiv. By far, the most preprints that were commented on, received only one comment (Fig. 4B).

**Figure 4.**
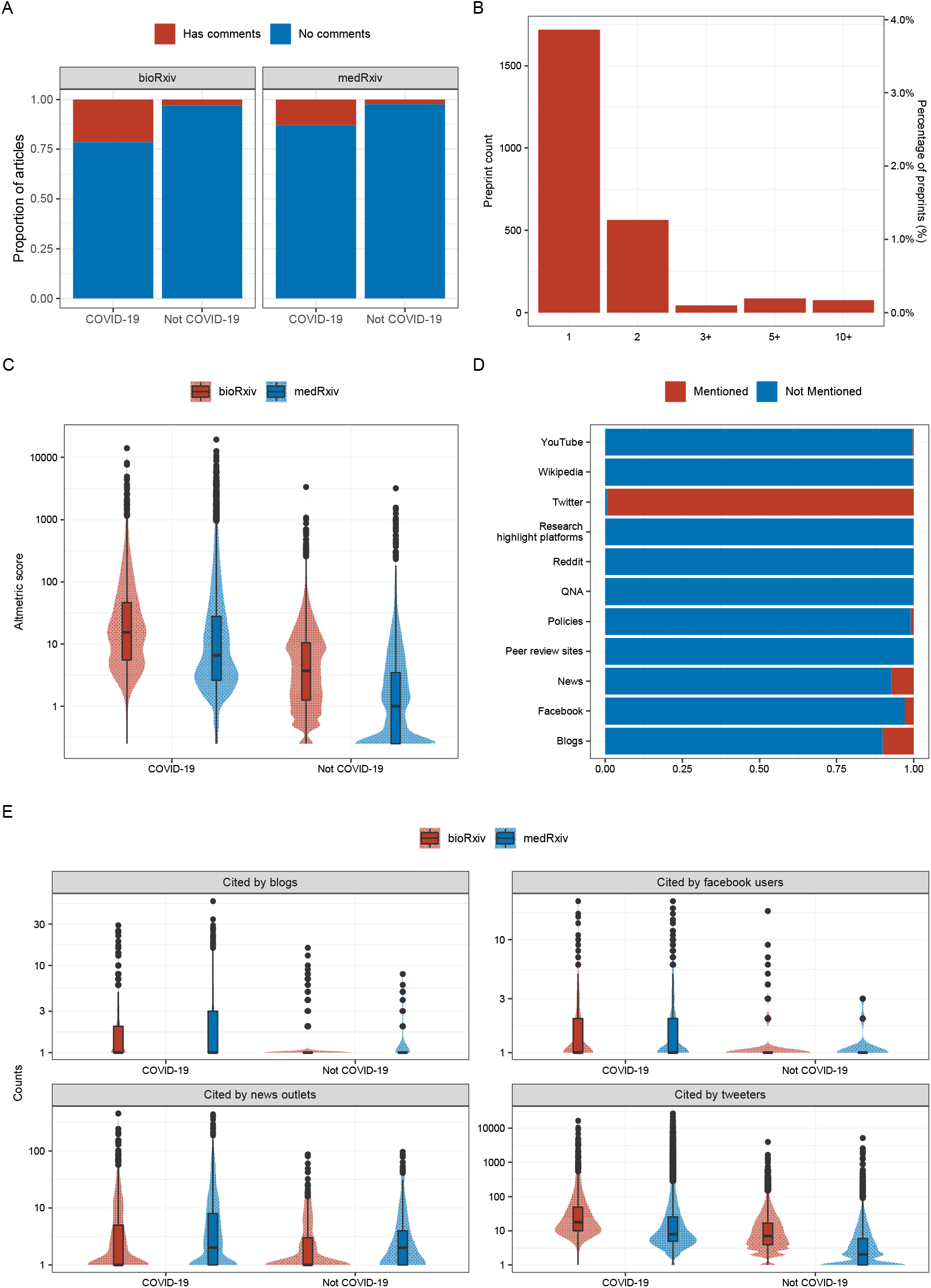
Disqus comments and Altmetric data for the identified preprints. **(A)** Percentage of preprints that received comments on bioRxiv or medRxiv website. **(B)** Distribution of comments number for preprints with comments. **(C)** Overall Altmetric score for all preprints on bioRxiv and medRxiv, with the first version released in 2020. **(D)** Percentage of preprints mentioned at various sources. **(E)** Counts of mentions by Facebook users, blog posts, news outlets, and Twitter. All data was retrieved on December 05-06, 2020.

Altmetric score, indicative of the public attention received by the preprints, appeared somewhat higher for COVID-19-related than for non-related preprints, particularly those posted on bioRxiv (vs. medRxiv) (Fig. 4C). Closer examination revealed that all preprints were mentioned on the Twitter platform (likely because of bots that tweet all deposited articles), a smaller percentage were mentioned in blog posts and news outlets, and a negligible number of preprints were mentioned on other venues (Fig. 4D). Grouped by the topic and preprint server, mentioning of the preprints on the Twitter platform closely reflected the overall Altmetric score. COVID-19-related preprints appeared more commonly shared on Facebook and blogs than non-COVID-19-related preprints (Fig. 4E). Preprints posted on the more clinically-oriented medRxiv were more reported in news outlets compared to those posted on bioRxiv (Fig. 4E).

### 3.3. Potential quality issues

A total of 345 notices were identified in the Retraction Watch Database, related to the articles published between January 01, 2020, and December 05, 2020, of which 46 were related to COVID-19. Comparison of the number of issued notices for COVID-19-related articles and articles pertaining to 4 different viruses and viral diseases and 2 different topics is depicted in Table 3. Briefly, only articles pertaining to herpes viruses had a higher approximated retraction rate than the COVID-19-related articles. COVID-19 articles had more retractions/expressions of concern/corrections than COVID-19-unrelated, articles pertaining to 3 viruses and their associated diseases (HIV, influenza, and hepatitis virus) and 2 topics related to COVID-19 (epidemiology of infectious diseases and immunology). Similarly, by searching PubMed we discovered that the retraction rate for COVID-19 articles was higher than for COVID-19-unrelated articles, although the difference is not so prominent (0.15‰ vs 0.13‰).

## 4. DISCUSSION

The present study was motivated primarily by previous observations of shorter submission-to-acceptance time for published COVID-19-related vs. non-related manuscripts (Homolak et al., 2020; Horbach, 2020; Kun, 2020), a phenomenon suggestive of preference of COVID-19-related manuscripts for publication in the peer-reviewed journals during the COVID-19 pandemic. The present data support such a view by demonstrating an independent association between COVID-19-related topic and a higher probability of publishing but have two major limitations that preclude straightforward generalizations: a) the analysis was limited only to a subset of all manuscripts “produced” during the observed period, i.e., those that were preprinted. This, however, was the only reasonable choice – these are the only manuscripts whose existence could be clearly verified and for which the risk (probability) of publishing could be (prospectively) estimated; b) the observed period was bounded (January 1 to November 01, 2020), which might have affected the outcomes: our supplemental analysis indicates that it could take up to around 500 days for a preprint to get published (Supplemental Figure S1), hence the present observations might simply reflect a certain lag-time present for non-COVID-19-related preprints. Therefore, the present results pertain and should be interpreted specifically with respect to preprinted manuscripts and the observed period which almost completely overlaps with the duration of the COVID-19 pandemic. We particularly accounted for potential bias arising from unequal “time at risk” by definition of two complementary outcomes differentially affected by the bounded observational period, and by stratification of preprints in respect to preprinting date. With adjustment for several other (albeit not all potentially relevant) potential sources of bias that could be captured in this kind of a study, the present estimates should be considered accurate. The observed considerably shorter submission-to-acceptance time for the published COVID-19-related vs. non-related preprints further supports the conclusion about the preference of COVID-19-related topics and is in line with observations pertaining to all published papers (Homolak et al., 2020; Horbach, 2020; Kun, 2020). In this respect, the present data should be viewed as reasonably indicative for all papers (during the COVID-19 pandemics) in general.

Increased preference (due to any reason) for the publishing of COVID-19-related (vs. non-related) papers combined with shorter submission-to-acceptance time, indicative of a shorter peer-review (and thus unlikely to be thorough and meaningful), creates a situation that could be reasonably considered susceptible to the impulsive release of publications of inadequate quality, i.e., susceptible to publishing bad or incorrect science, or just nonsense (e.g. article reporting a link between 5G and SARS-CoV-2) (Fioranelli et al., 2020). At least theoretically, preprinting provides a (possible) way to ameliorate this problem by opening a time window for public pre-submission peer-review that could complement the journal peer-review. The extent, quality and relevance of any peer-review, and in particular the public pre-submission peer-review, is difficult to quantify. The present data, using the number of comments pertaining to preprinted manuscripts as a proxy, do not suggest that such a practice is actually common: only around 5% of the preprints were commented on, typically with only one comment. On a positive side, COVID-19-related papers received more comments (than non-related), suggesting that these preprints are (at least) publicly discussed. Just as observed by others (Yeo-Teh and Tang, 2020), the present results indicate that published COVID-19 articles, at the present state, have a higher retraction rate than non-COVID-19 articles. However, our data should be viewed with additional caution – reporting retractions, corrections, and expression of concern as fractions with numbers obtained from PubMed is not optimal and is only an approximation. Furthermore, as Abritis et al. have warned (Abritis et al., 2020), it is hard to compare the number of retracted articles given that it takes years for article retractions. There is a possibility that the retraction numbers of non-COVID-19 articles are only lagging and will eventually catch up with the COVID-19 articles. Additionally, higher retraction rates might reflect greater public scrutiny, not necessarily lower quality.

While not the primary focus of this article, our limited data supports claims that preprinting in the biomedical field has increased during the COVID-19 pandemic (Fraser et al., 2020). Several other preprint-related activities, like deposition of more than one preprint version, shortened time between versions, or changes of preprint titles (as compared to the time period before the pandemic) also seem to be intensified (see Supplemental Figure S2 and S3). We hold this fact to be much needed and long overdue. We believe that unfeigned quality concerns due to lack of peer-review are (more or less) surmountable by the audiences’ critical approach. On the other hand, lack of peer-review and editorial screening might be advantageous with respect to the speed of information sharing, open discussion, and lack of “censorship”. This could reduce the need for hasty publication of inadequate papers in scientific journals. In addition to causing a distrust towards science, the damage done by the poor journal-published articles is hard to rectify, largely due to the false sense of unquestionable credibility assigned to articles published in peer-reviewed journals. This concern in the context of COVID-19 pandemic has already been recognized and brought up by Serge Horbach declaring that “nonsense or incorrect science in one of these papers is potentially much more harmful” (Kwon, 2020). One illustrative example from the past is the infamous case of the article reporting a link between MMR vaccine and autism that drives distrust towards vaccination even today, years after its retraction (Omer, 2020).

Finally, in addition to the advantages for the profession and science, preprints might be a valuable “tool” for researching science itself. Preprinting provides an opportunity to study what happens “behind the curtains” of the hidden journal submission process (i.e. provides an opportunity to study the peer-review itself). We hope this will be recognized and utilized in the future. However, incorrect identification of the journal published preprints is one of the obstacles that have yet to be overcome. A significant number of preprints has not been identified as published by bioRxiv/medRxiv services (Abdill and Blekhman, 2019), even though the declaration is posted on their website stating that this happens “on rare occasions” because authors or titles have changed (“Frequently Asked Questions (FAQ),” n.d.). This is potentially (if the error rate is not similar in COVID-19-related vs -unrelated preprints) the most significant limitation of this study.

In conclusion, during the COVID-19 pandemic, there appeared an increasing preprinting trend on bioRxiv or medRxiv. In the case of the latter platform, the trend is primarily due to the preprinting of COVID-19-related manuscripts. COVID-19-related preprints are more likely to be published in the peer-reviewed journals and their submission-to-acceptance time (a proxy for the peer-review process) is considerably shorter than for the COVID-19 non-related manuscripts. COVID-19-related preprints received more comments on the preprinting platforms, but the proportion of preprints commented-on is generally modest. This suggests that the opportunity of public pre-submission peer-review, inherent to the concept of preprinting, is not seized to any relevant extent. Retractions and issued concerns/corrections were sporadic regarding the papers published (and indexed in PubMed) between January 1 and December 05, 2020, but the incidence of retractions/concerns/corrections was higher for published COVID-19-related than for non-related papers. Overall data support a view that so far, COVID-19-related manuscripts were favored for publishing in peer-reviewed journals, commonly with very short peer-review.

## Acknowledgments

Nothing to acknowledge.

## 5. DECLARATIONS

### Ethics approval

None applicable.

### Authors’ contributions

IK and VT designed the study. IK gathered data. IK and VT analyzed the data. DV and JH provided valuable suggestions for analyzing data. All authors participated in the writing of the manuscript.

### Data availability

Data and *R* code is available on GitHub (ikodvanj/publishing_of_preprints).

