## Supplemental for "Publishing of COVID-19 Preprints in Peer-reviewed Journals, Preprinting Trends, Public Discussion and Quality Issues"

### Supplemental Material

To supplement the observations of the main study, we evaluated preprinting trends on bioRxiv and medRxiv since their start, as well as trends in probability of publishing of preprinted manuscripts and trends in the deposition of preprint versions.

### Supplemental methods

Essentially the same methodology was used just as for the main study. *Rbiorxiv* and *medrxivr* R packages were used to access bioRxiv and medRxiv application programming interfaces (APIs) and to collect metadata for all preprints deposited on bioRxiv (since 2013) and medRxiv (since 2019). COVID-19 preprints were identified using the search terms COVID-19 OR SARS-CoV-2 OR Coronavirus disease 19 OR 2019-nCoV. All other preprints were classified as non-COVID-19 preprints. Publication dates of articles that were preprinted on bioRxiv were provided by the bioRxiv API. *Rcrossref* was used to gather publication dates for journal-published preprints initially deposited on the *medRxiv*.

Time-to-publishing was summarized as Kaplan-Meier curves by preprint server and 6 months intervals (based on the date of the first deposited preprint version).

All other metadata, including the number of preprint versions, the time difference between versions and changes in titles of preprints, were summarized descriptively.

### Supplemental results

Considering the entire time period of existence of bioRxiv and medRxiv, approximately 65-70% of all preprinted articles eventually got published. The lag time between deposition of the first preprint version and publishing can extend to up to several hundreds of days (Figure S1).

A difference in the maximum number of versions can be observed for preprints deposited during 2020 as compared to previous years. COVID-19-related and subsequently published preprints seem to have more deposited versions than COVID-19 preprints not published by the cut-off date (November 01, 2020) (Figure S2A). This trend that (eventually) published preprints have more deposited versions can also be observed in previous years (Figure S2B). The number of days elapsed between preprint versions seems to be similar for COVID-19 published and (still) not published preprints. On the other hand, COVID-19 unrelated published preprints seem to have lengthier time between versions than unpublished - a pattern that can be observed to some extent also for preprints deposited during previous years (Figure S2C-D).

The percentage of preprints that changed title between versions seems to be increasing over the years (Figure S3A). This might suggest that during previous years, preprints were deposited as finished works, with only minor changes introduced to subsequent versions (if any). On the other hand, it seems that the perception of preprints has been changing and that preprints are more and more being posted as “works in progress”. Considering only the year 2020, the percentage of preprints with changes in titles between versions is higher for COVID-19 related than non-related preprints (Figure S3B).

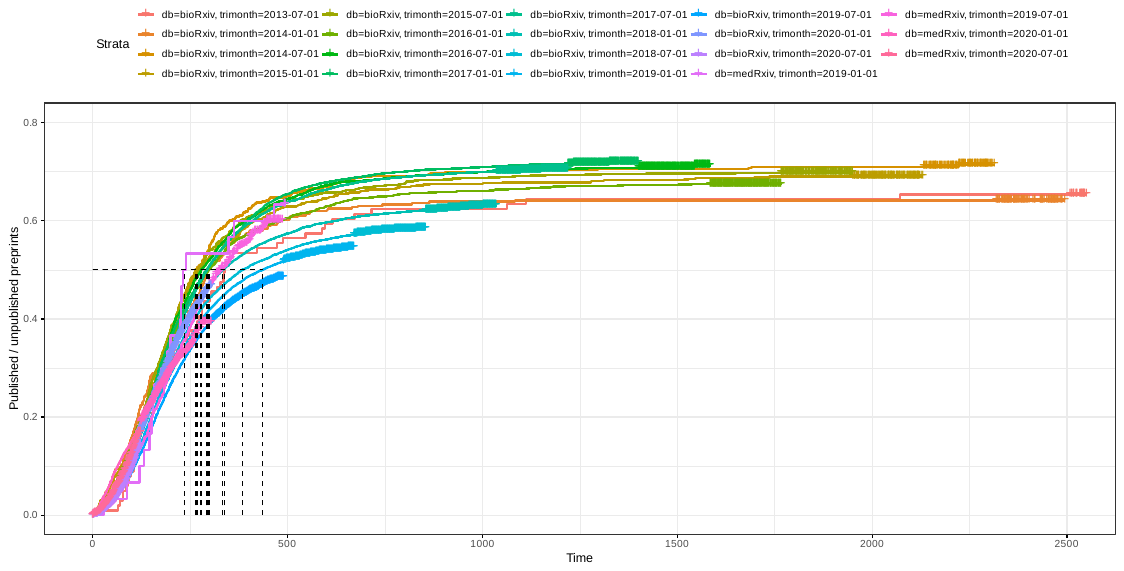

**Figure S1.** Kaplan-Meier curves summarizing time-to-publishing (cut-off date Nov. 01, 2020) considering all preprints deposited on bioRxiv and medRxiv since their inception, stratified by the date of deposition of the first version (6-month interval strata).

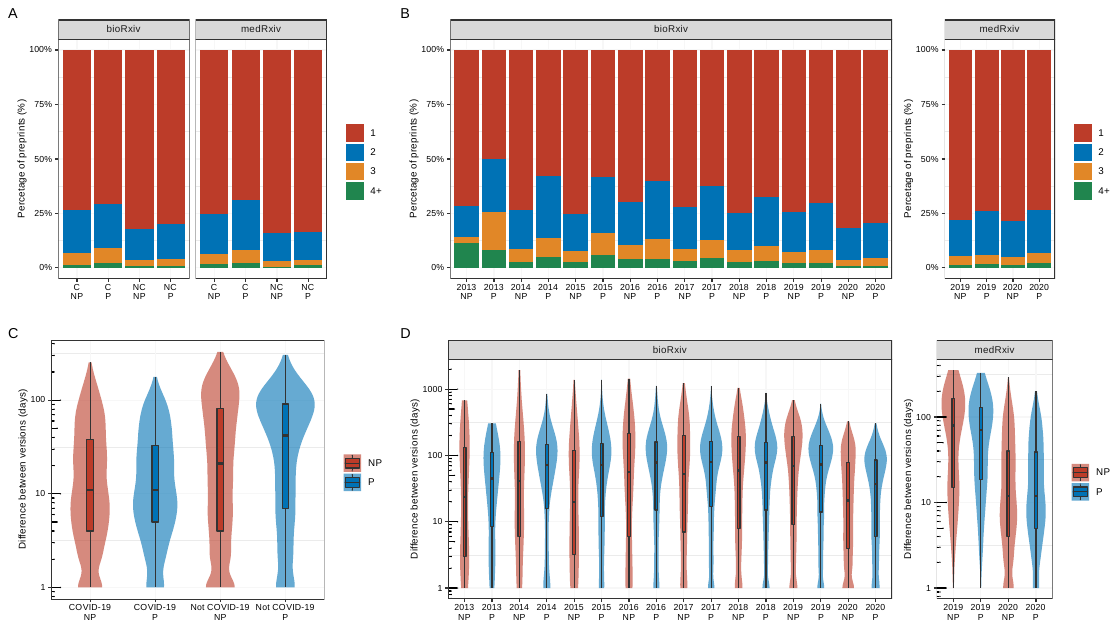

**Figure S2.** The proportion of preprints with only one (final) or more versions (1, 2, 3 or 4+) and time interval (in days) between versions**.** A and C display data pertaining to preprints deposited during 2020, B and D display data on preprints deposited during preceding years. *C* - COVID-19, *NC* - not-COVID-19, *NP* - not published, *P* - published.

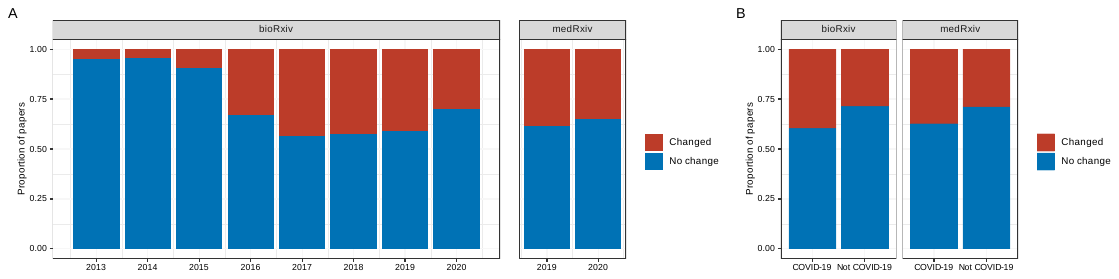

**Figure S3.** Changing of the article title between preprint versions. (A) The proportion of articles that changed the title at least once between versions, with respect to the preprinting repository and the publication year of the first version. (B) The proportion of preprints deposited during 2020 on bioRxiv and medRxiv that changed the title at least once between versions.
